# Comparative study of four SARS-CoV-2 Nucleic Acid Amplification Test (NAAT) platforms demonstrates that ID NOW performance is impaired substantially by patient and specimen type

**DOI:** 10.1101/2020.06.04.135616

**Authors:** Paul R. Lephart, Michael Bachman, William LeBar, Scott McClellan, Karen Barron, Lee Schroeder, Duane W. Newton

## Abstract

The advent of the COVID-19 pandemic in the United States created a unique situation where multiple molecular diagnostic assays with various indications for use in the detection of SARS-CoV-2 rapidly received Emergency Use Authorization by the FDA, were validated by laboratories and utilized clinically, all within a period of a few weeks. We compared the performance of four of these assays that were being evaluated for use at our institution: Abbott RealTime m2000 SARS-CoV-2 Assay, DiaSorin Simplexa COVID-19 Direct, Cepheid Xpert Xpress SARS-CoV-2 and Abbott ID NOW COVID-19. Nasopharyngeal and nasal specimens were collected from 88 ED and hospital-admitted patients and tested by the four methods in parallel to compare performance. ID NOW performance stood out as significantly worse than the other three assays despite demonstrating comparable analytic sensitivity. Further study determined that the use of a foam nasal swab compared to a nylon flocked nasopharyngeal swab, as well as use in a population chronically vs. acutely positive for SARS-CoV-2, were significant factors in the poor comparable performance.

## Introduction

The rapid onset of COVID-19 in the United States resulted in an accelerated pace of both SARS-CoV-2 nucleic acid amplification test (NAAT) development and FDA Emergency Use Authorization (EUA) approvals. Clinical microbiology laboratories that typically would take weeks to evaluate and verify performance characteristics for a FDA approved diagnostic test had little choice but to perform abbreviated validation and/or verification studies of assays, benchmarked against limited FDA EUA performance data, in a matter of days. The sheer volume of COVID-19 test requests from different patient populations, different specimen types, and with different turnaround time needs demanded that laboratories implement more than one type of NAAT to respond to the crisis.

SARS-CoV-2 testing in our laboratory began with the CDC EUA assay performed on Abbott m2000, but due to that assay’s significant throughput constraints (24 specimens in 8 hours), we quickly verified and switched to the Abbott RealTime SARS-CoV-2 EUA Assay (m2000) once released (94 specimens in ~8 hours). Although this assay provided capacity for our outpatient testing needs, we also verified the DiaSorin Simplexa COVID-19 Direct (Simplexa) assay, capable of resulting 8 specimens in 90 minutes, and used this assay as a rapid turn-around time (TAT) option for our inpatient and emergency department (ED) populations. Within a few weeks, additional SARS-CoV-2 NAAT options emerged that were specifically designed for rapid testing of patients in the point of care setting: the Cepheid Xpert Xpress SARS-CoV-2 (Xpert) assay, which could provide results in 45 minutes, and the Abbott ID NOW COVID-19 (ID NOW) assay, ultimately approved for direct nasal, nasopharyngeal and throat swab testing only, with results in 5-15 minutes.

In the absence of clinical trials and a gold standard for COVID-19 diagnosis, the clinical performance of SARS-CoV-2 assays is unclear. Anecdotal claims of poor NAAT performance exist in the lay press, and limited studies have shown variable performance of rapid POC tests (1–9). As a surrogate for a gold standard, a composite reference standard (CRS) can be used to determine the consensus of comparable assays and identify outlier assays in terms of clinical performance (10). Using this approach, our goal was to evaluate the performance—in parallel— of three NAATs from nasopharyngeal (NP) swabs in M4-RT viral transport medium (VTM) (m2000, Xpert, Simplexa) and an NAAT assay performed directly from a nasal swab (ID NOW).

## Methods

### Accuracy Study Design

From 4/22 – 5/5/2020, specimens were collected from 88 ED and hospital admitted patients and tested for SARS-CoV-2 on the RealTime m2000 SARS-CoV-2 Assay (Abbott Molecular, Des Plaines, IL), Simplexa™ COVID-19 Direct (DiaSorin, Cypress, CA), Xpert^®^ Xpress SARS-CoV-2 (Cepheid, Sunnyvale, CA) and ID NOW COVID-19 (Abbott Molecular, Des Plaines, IL) within 24 hours of collection. Each assay was performed according to manufacturer’s EUA instructions.

NP and nasal swabs were collected from 88 patients, of which 75 were patients presenting in the ED and 13 were from a population of recovering COVID-positive inpatients. NP specimen collection, transport to the hospital-based core microbiology laboratory in M4-RT viral transport medium (Thermo Fisher Scientific, San Diego, CA; VTM) and subsequent testing by Simplexa was performed as a part of routine clinical care. Residual NP specimen in VTM was stored at 4°C, transported to our offsite main laboratory, and within 24 hours of collection, used for comparative study testing by m2000 and Xpert assays. At the time of NP swab collection, nasal swabs were collected in parallel from each of these patients and transported dry to the offsite main laboratory in a sealed sterile collection bag, stored at 4°C and tested by ID NOW within 24 hours, consistent with the package insert procedure.

In order to determine a percent agreement amongst the methods, we established a composite reference standard as defined by result agreement of SARS-CoV-2 target in at least 2 of 4 NAAT results. Agreement for each individual assay was compared to this standard.

### Analytic Limit of Detection Study Design

Dilutions were prepared from ZeptoMetrix inactivated SARS-CoV-2 virus (1.70 × 10^5^ TCID50/ml) that was internally quantified to 10^9.62^ copies/mL relative to a standard curve of AccuPlex™ SARS-CoV-2 Reference Material (SeraCare) constructed on the m2000 assay. From that stock, dilutions were made in VTM (1,042, 521, 260, 130, 65 and 32.5 copies/mL) and tested on the m2000, Simplexa, and Xpert SARS-CoV-2 assays. A separate set of ZeptoMetrix dilutions in VTM were made so that when added to the 2.5 ml of elution buffer in the ID NOW receiver cup, equivalent concentrations were achieved (1,042, 522, 262, 105 and 53 copies/mL). In this way, the concentrations of viral copies in the ID NOW buffer were equivalent to the VTM dilutions used for the other three instruments. Five replicates at each dilution were tested.

## Results & Discussion

Nasal swabs directly tested on the ID NOW assay had 48% positive agreement compared to the CRS, whereas Simplexa had 88%, m2000 had 96% and Xpert had 100% positive agreement (Table 1). While the deficit in positive percent agreement (PPA) seen in ID NOW test results is consistent with other early release studies in the scientific literature ((1–4, 7–9)), it is surprising given the ID NOW’s LOD claim of 125 genome equivalents/mL, which is similar to the 100 copies/mL claimed by the m2000 method, 250 copies/mL claimed by Xpress and 242 copies/mL claimed by Simplexa. To clarify this apparent discrepancy, a direct assessment of the analytic sensitivity of all assays in this comparison was performed utilizing dilutions of inactivated SARS-CoV-2 whole virus (ZeptoMetrix). In this limited LOD study, we found each assay had comparable LODs to the reported LOD data in their package insert, including the LOD of the ID NOW assay, as shown in Table 2.

**Table 1a:** Agreement of Four SARS-CoV-2 NAATs relative to the CRS

|  | Composite Reference Standard (CRS) |  | Percent Agreement with CRS |  | 95% CI |
| --- | --- | --- | --- | --- | --- |
|  | Detected | Not detected |  |  |  |
| **ID NOW** |  |  | Positive Agreement = | 48% | 0.30 to 0.67 |
| Detected | 12 | 0 | Negative Agreement = | 100% | 0.94 to 1.0 |
| Not detected | 13 | 63 | Overall Agreement = | 85% |  |
| **Simplexa** |  |  | Positive Agreement = | 88% | 0.70 to 0.96 |
| Detected | 22 | 0 | Negative Agreement = | 100% | 0.94 to 1.0 |
| Not detected | 3 | 63 | Overall Agreement = | 97% |  |
| **m2000** |  |  | Positive Agreement = | 96% | 0.80 to 1.0 |
| Detected | 24 | 0 | Negative Agreement = | 100% | 0.94 to 1.0 |
| Not detected | 1 | 61 | Overall Agreement = | 99% |  |
|  |  | 2 invalids |  |  |  |
| **Xpert** |  |  | Positive Agreement = | 100% | 0.87 to 1.0 |
| Detected | 25 | 2 | Negative Agreement = | 97% | 0.87 to 0.99 |
| Not detected | 0 | 60 | Overall Agreement = | 98% |  |
|  |  | 1 invalid |  |  |  |

**Table 2:**

|  | Study LOD in M4-RT<br>(copies/ml)* | Package insert LOD | Average Ct at LOD |
| --- | --- | --- | --- |
| m2000 | 32.5 | 100 copies/ml | 26.5** |
| Cepheid | 65 | 250 copies/ml | 36.7 / 39.8 |
| ID NOW | 262 | 125 genome equivalents/ml | N/A |
| Simplexa | 521 | 242 copies/ml | 32.6 / 33.0 |

\* Defined as lowest dilution in which 5/5 replicates detected.
\*\* Reported Ct value for m2000 excludes 10 unread cycles.

Similarities in the LODs among the assays suggest that other factors contribute to the differences in comparative performance when testing clinical specimens. When additional ID NOW testing was performed on the 25 NP VTM specimens that were positive based on the CRS, 6 additional patients were detected that were negative by nasal swab testing. This suggests that testing an NP swab in VTM on ID Now may have better performance than a direct nasal swab, as use of the NP VTM improves the PPA from 48% to 64%. However, testing an NP swab in VTM was recently removed as an approved source from the original ID NOW package insert based on concerns about false negatives, whereas a nasal swab is provided as an approved collection device. The sensitivity issues leading up to that removal may have less been due to VTM than the quality of a nasal specimen itself when compared to a NP specimen. Further studies should be conducted directly comparing the performance of direct (not placed in VTM) NP swabs on the ID NOW to NP swabs in VTM tested by other NAATs.

An important variable in our study is that two distinct population groups were analyzed. Thirteen of the 88 patient specimens collected were from an inpatient population of recovering COVID positive patients, with a mean time from initial COVID-19 diagnosis of 25.8 days. A comparison of the m2000 Ct values obtained from m2000 positive samples from inpatients (red) and ED patients (black) and by ID NOW result is shown in Figure 1. The mean m2000 Ct value for the ID NOW positive specimens was 14.3 versus a significantly higher mean m2000 Ct value 22.29 (p-value < 0.001) for the ID NOW negative specimens. Inpatients had significantly lower Ct values as measured by Abbott m2000 Ct (n=9, mean Ct 21.6) than positive specimens collected from patients presenting at the ED (n=16, mean Ct 16.3, p-value 0.04). As the majority of ID NOW negative/m2000 positive specimens (8 of 12) were from this inpatient population of low Ct positives, the overall performance of the ID NOW assay was substantially impacted. To assess test performance in a more typical use case in a POC setting, we reanalyzed percent agreement for all assays using only ED patients. While still notably lower than the other assays, the PPA of ID NOW increased from 48 to 69%, whereas performance of the other assays was nearly identical (Table 1b).

**Figure 1:**
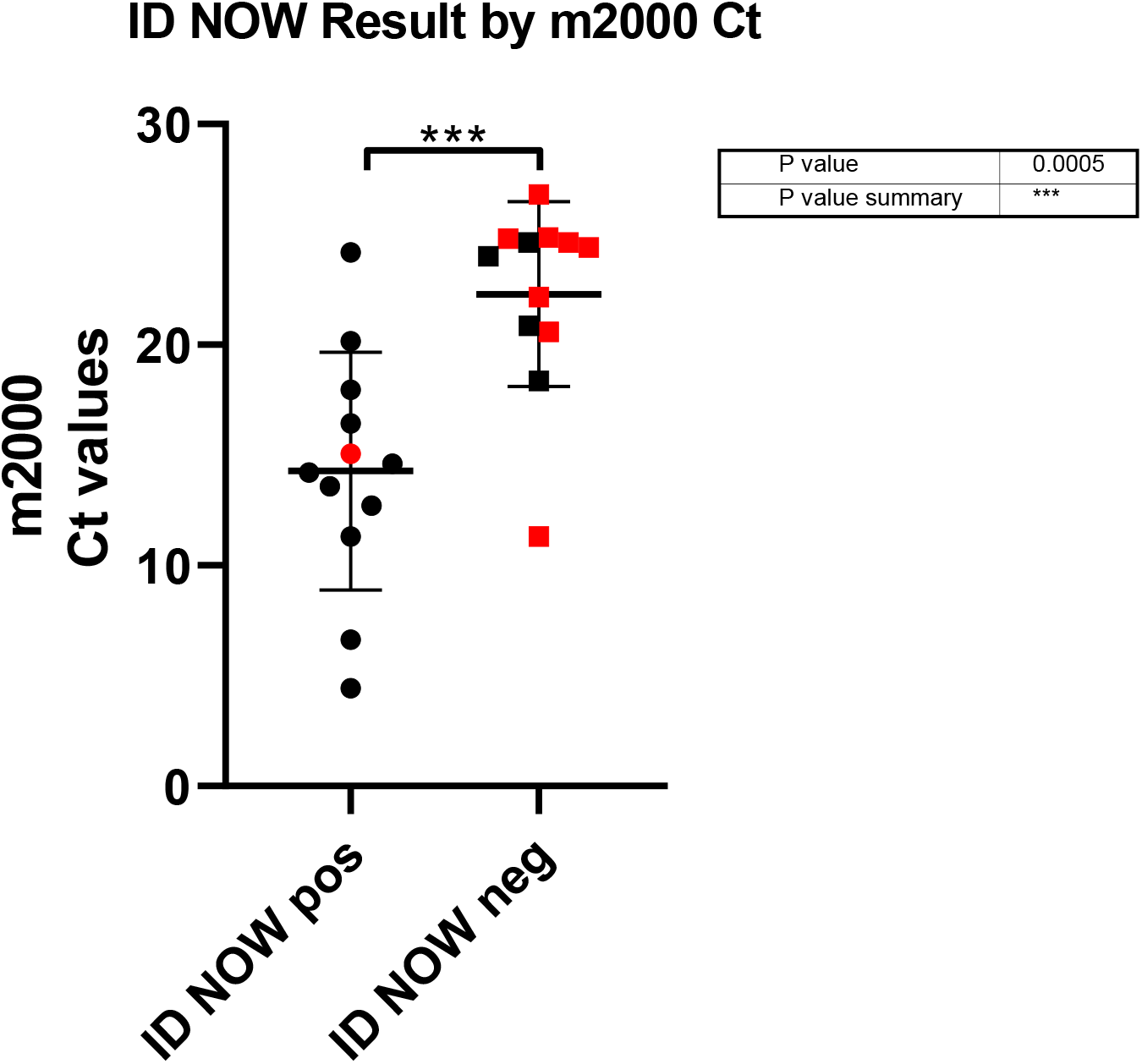
ID NOW Result by m2000 cycle time. Data points depicted in red indicate inpatient specimens and black are ED specimens.

**Table 1b:** Agreement of Four SARS-CoV-2 NAATs relative to the CRS (ED patients only)

|  | Composite Reference Standard (CRS) |  | Percent Agreement with CRS |  | 95% CI |
| --- | --- | --- | --- | --- | --- |
|  | Detected | Not detected |  |  |  |
| **ID NOW** |  |  | Positive Agreement = | 69% | 0.44 to 0.86 |
| Detected | 11 | 0 | Negative Agreement = | 100% | 0.94 to 1.0 |
| Not detected | 5 | 59 | Overall Agreement = | 93% |  |
| **Simplexa** |  |  | Positive Agreement = | 88% | 0.64 to 0.98 |
| Detected | 14 | 0 | Negative Agreement = | 100% | 0.94 to 1.0 |
| Not detected | 2 | 59 | Overall Agreement = | 97% |  |
| **m2000** |  |  | Positive Agreement = | 94% | 0.72 to 1.0 |
| Detected | 15 | 0 | Negative Agreement = | 100% | 0.94 to 1.0 |
| Not detected | 1 | 57 | Overall Agreement = | 99% |  |
|  |  | 2 invalids |  |  |  |
| **Xpert** |  |  | Positive Agreement = | 100% | 0.81 to 1.0 |
| Detected | 16 | 2 | Negative Agreement = | 97% | 0.89 to 0.99 |
| Not detected | 0 | 58 | Overall Agreement = | 98% |  |
|  |  | 1 invalid |  |  |  |

This comparative analysis of SARS-CoV-2 NAATs utilizing the m2000, Simplexa, Xpert and ID NOW assays demonstrated that significant performance deficits were found in the ID NOW assay when tested in a mixed patient population using both NP and nasal specimens. Based on a CRS, use of the m2000, Xpert, and Simplexa assays for NP specimens in VTM are likely to have similar performance in clinical practice and choice of implementation can be made based on considerations of turnaround time, throughput, work flow and cost. In contrast, despite the ID NOW assay claiming and demonstrating comparable (differences < 1 log_10_) analytic LOD findings to the other assays tested, the lower detection rate of the ID NOW from nasal samples must be considered when deciding on a use case. When limiting our data set to an acute ED patient population and comparing results from the same specimen type (NP in VTM), performance of ID NOW was improved but still demonstrated lower performance compared to the other assays tested. In situations where the 5-15 minute turn-around time of the ID NOW provides distinct advantages, it is critical that the most appropriate specimen type, appropriate patient population and need for more sensitive confirmatory NAAT testing be assessed prior to use.

